# Protein-Protein interaction and quantitative phosphoproteomic studies reveal potential mitochondrial substrates of protein phosphatase 2A-B’ζ holoenzyme

**DOI:** 10.1101/2023.05.20.541568

**Authors:** Ahmed Elshobaky, Cathrine Lillo, Kristian Persson Hodén, Amr R.A. Kataya

## Abstract

Protein phosphatase 2A (PP2A) is a heterotrimeric conserved serine/threonine phosphatase complex including a catalytic, scaffolding, and regulatory subunit. Three A Subunits, 17 B subunits, and five C subunits are encoded by the *Arabidopsis* genome, allowing 255 possible PP2A holoenzyme combinations. The regulatory subunits are crucial for substrate specificity and PP2A complex localization and are classified into B, B’, and B’’ non-related families in land plants. In *Arabidopsis*, the close homologs B’η, B’θ, B’γ, and B’ζ are further classified into a subfamily of B’ called B’η. Previous studies suggest a role of mitochondrial targeted PP2A subunit (B’ζ) in energy metabolism and plant innate immunity. Potentially, the PP2A-B’ζ holoenzyme is involved in the regulation of the mitochondrial succinate/fumarate translocator or affects enzymes involved in energy metabolism. To investigate this hypothesis, the interaction between PP2A-B’ζ and enzymes involved in the mitochondrial energy flow was investigated using bimolecular fluorescence complementation in tobacco and onion cells. Interaction of B’ζ subunit was confirmed with the Krebs cycle proteins Succinate/fumarate translocator (mSFC1), Malate dehydrogenase (mMDH2), and Aconitase (ACO3). Additional putative interacting candidates were deduced from comparing the enriched phosphoproteomes of wild type and B’ζ mutants: the mitochondrial regulator *Arabidopsis* pentatricopeptide repeat 6 (PPR6) and the two metabolic enzymes Phosphoenolpyruvate carboxylase (PPC3) and Phosphoenolpyruvate carboxykinase (PCK1). Overall, this study identifies potential PP2A substrates and highlights the role of PP2A in regulating energy metabolism in mitochondria.

## Introduction

Protein post-translational modifications (PTMs) are covalent additions or modifications of chemical groups to a protein after its synthesis. PTMs add an additional level of regulation, on top of gene expression and mRNA translation. Phosphorylation is an abundant PTM and a main regulator of cell signaling that is involved in a vast number of cellular processes [1]. Reversible phosphorylation is achieved through the balance between protein kinases, which act by adding a phosphate group to amino acids in the protein chain, and phosphatases, which remove phosphorylated groups from proteins [2]. Attesting to the importance of protein phosphorylation events, the *Arabidopsis thaliana* genome contains around 940 kinases and 150 phosphatases [3]. Protein phosphorylation and dephosphorylation processes regulate multiple downstream cellular processes such as, e.g., transcription, energy metabolism and development, by affecting protein attributes such as, e.g., localization, interactions, and activity [1,3].

Meta-analysis of phosphoproteomics data revealed that serine (Ser) is the most abundantly phosphorylated residue in plants (∼85%), followed by threonine (Thr; ∼15%) [4]. Consequently, one of the major phosphatase families in plants is composed of the serine/threonine-specific phosphoprotein phosphatases (PPPs). Among the PPPs, PP1 and PP2A are the largest groups in *Arabidopsis* [5,2]. Plant PP2A enzymes have been shown to be involved in various functions such as regulation of light signaling, hormone signaling, metabolism, other PTMs and defense signaling [5,6]. PP2A enzymes are heterotrimeric proteins composed of a catalytic C subunit with regulatory (B) and scaffolding (A) subunits. The regulatory subunits are essential for substrate specificity and localization of the complex. Complexity of PP2A enzymes is achieved through combination of a low number of C subunits with A and B subunits [7]. Thus, despite having only five C subunits, *Arabidopsis* can theoretically attain over 250 PP2A variants (three A subunits and 17 B subunits). The plant B family is further divided into the B, B’ and B’’ subfamilies, of which the B’ gene family contains nine members: α, β, γ, δ, ε, ζ, η, θ and κ [2]. We here focus on the B’ subunit ζ (Z).

The plant immune system needs to be under tight negative regulation to avoid over-activation and autoimmunity. Protein kinases and phosphatases have been suggested as key players in plant immunity regulation, and some emergent studies have implicated PP2A in this role [8,9,10]. More specifically, all four members of the B’η subgroup (η, γ, θ and ζ) were shown to be negative regulators of plant PAMP-triggered immunity. Increased resistance to the virulent bacterial strain *Pseudomonas syringae* DC3000 was observed in knockout mutants of *b’θ, b’ζ* and *b’η*. Furthermore, the PP2A-B’ζ and PP2A-B’η complexes were found to control the activation of cell-surface pattern recognition receptors by regulating the phospho-status of the co-receptor BAK1 [8]. The *b’γ* knockdown, on the other hand, displayed constitutive defense responses, and, in line with this, was implicated in resistance to a variety of infectious agents; aphids, hemibiotrophic bacteria and necrotrophic fungi [10,11].

Notably, the constitutive defense activation observed in the *b’γ* mutant seems to be explained by an involvement of the PP2A-B’γ complex in methionine recycling. By performing bimolecular fluorescence complementation (BiFC) experiments, Rahikainen et al. [12] first showed that B’γ interacts with indole glucosinolate methyl transferases. Further, B’γ was found to negatively regulate the metabolism of glucosinolates, which are important secondary metabolites, acting as repellents for aphids and microbial plant pathogens in *Arabidopsis*. Increased build-up of specific compounds such as 4-methoxy-indol-3-yl-methyl glucosinolate is therefore a likely reason for the elevated defense responses of the *b’γ* knockout. All of which solidify the importance of the B’η subgroup in plant immune responses.

Mitochondria are double membrane-bound eukaryotic organelles. Reversible phosphorylation involving mitochondrial phosphatases has been largely neglected, although mitochondria are increasingly recognized as a hub for cell signaling, and several mitochondrial phosphatases have been reported to play vital functions. Nonetheless, evidence for the localization and activities of these reported phosphatases, as well as their functions and their mitochondrial translocation mechanism, are still poorly understood [13].

PP2A regulatory subunit (B’ζ) was reported to target mitochondria [7], and when B’ζ is mutated, plants not only show negative regulation of plant innate immunity, but also have a sugar dependence phenotype [8,9]. This indicates a defect in energy metabolism in the *pp2a-b’ζ* mutant. As seedlings of the knockout mutant of PP2A-B’θ, the nearest homolog of PP2a-B’ζ, have shown a defect in peroxisomal β-oxidation, and is assumed to block the supply of succinate to the mitochondrial Krebs cycle during early seedling establishment [14], we investigated a putative role for PP2A-B’ζ in the regulation of energy flow in mitochondria. We here studied *in vivo* interactions with proteins involved in the energy flow to mitochondria and identified potential substrates of PP2A-B’ζ employing protein-protein interaction and comparative quantitative phosphoproteomics.

## Methodology

### Plant material and growth conditions

Seeds of *Arabidopsis thaliana* wild type (Col-0) and two *pp2a-b’ζ* knockout lines (SALK_150586C; “*z1*” and SALK_107944C; “*z2*”) were surface-sterilized and stratified. Plants were grown under short day or long day conditions as described in [15]. Mutant seeds were obtained from the European Arabidopsis Stock Centre (Nottingham, UK) and genotyped by PCR using T-DNA insertion primers obtained from SALK institute website SIGnAL (http://www.signal.salk.edu/tdnaprimers.2.html) as shown in Supplementary Table S1.

### Gene cloning for *in planta* expression *in vivo*

Transient expression of constructs for BiFC was done in two systems: *Agrobacterium* infiltration into *Nicothiana benthamiana* leaves and gene gun delivery into onion epidermal cells. BiFC constructs for onion transformation were created by cloning the genes for candidate interactors into pUC-VYNE_N173_ and the gene for B’ζ (At2g21650) into pUC-VYCE_C155_. For *Agrobacterium* constructs, the plasmids hygII-VYNE_N173_ and kanII-VYCE_C155_ were used for interactors and B’ζ, respectively (Supplementary Fig. S1). For *mMDH2* and *CSY5*, RNA was extracted from *Arabidopsis* flower tissue (Qiagen RNeasy Plant Mini, 74004), reverse transcribed using SuperScript IV (ThermoFisher, 18090010) and PCR-amplified using Phusion polymerase (ThermoFisher, F531L). Genes from rest of the interactor candidates were obtained in plasmids from ABRC (https://abrc.osu.edu). Correctness of the inserts was verified by Sanger sequencing. All cloning primers are listed in Supplementary Table S1.

*N. benthamiana* plants were grown in soil under long day conditions and transformed as described in [16]. Gene gun transformation into onion epidermal cells was done as in [15]. Interactions were analysed two days post infiltration for tobacco leaves and one to two days after bombardment for onion cells as described by [14]. Each interaction was analysed in at least three independent experiments.

### Sugar dependence assay

Seedlings were grown on Linsmeier and Skoog (LS) for 7 days with or without sucrose (1%) under short-day conditions (light:dark 8h:16h) or in continuous darkness. The seedlings were scanned using a CANON scanner, and hypocotyl length was measured using ImageJ (https://rsb.info.nih.gov/ij/).

### Quantitative phosphoproteomics

Total protein was extracted from 170 mg tissue collected from 6-day-old seedlings grown on Murashige and Skoog (MS) without sugar (triplicate samples each from WT, *z1* and *z2*) as described by Roitinger et al. [17]. The proteins were precipitated using the TCA/acetone method, collected by centrifugation, and processed for trypsin digestion using filter-aided sample preparation. Digestion was done overnight at 37°C using 0.22 µg/µL MS-grade trypsin (Promega, PRV5111) and the samples were purified on peptide desalting spin columns (Thermo Fisher Scientific, 89851). The phosphopeptides were labeled with nine isobaric tandem mass tags, using the TMT10plex kit (Thermo Fisher Scientific, 90110), and enriched by Sequential enrichment of Metal Oxide Affinity Chromatography (High-Select SMOAC; Thermo Fisher Scientific, A32993).

### Liquid chromatography-tandem mass spectrometry (LC-MS/MS)

Tryptic peptides were analysed by LC-MS/MS at the Southern Alberta Mass Spectrometry Facility using an Orbitrap Fusion Lumos Tribrid mass spectrometer (Thermo Fisher Scientific) controlled by Xcalibur (v4.0.21.10) and connected to a Thermo Scientific Easy-nLC 1200 system. Briefly, the Orbitrap first performed a full MS scan at a resolution of 120000 FWHM to detect the precursor ion (m/z from 375 to 1575, charge of +2 to +4). The AGC (auto gain control) and the max injection time were respectively set at 4e5 and 50 ms. The most intense precursor ions that showed a peptidic isotopic profile and had an intensity threshold over 2e4 were isolated using the quadrupole (window 0.7) and fragmented using HCD (38% collision energy) in the ion routing multipole. Fragment ions (MS2) were analyzed at a resolution of 15000, setting the first mass to 100 in order to acquire the TMT reporter ions (AGC 1e5 ms, max injection time 105 ms). To avoid acquisition of the same precursor ion having a similar m/z (plus or minus 10 ppm), dynamic exclusion was enabled for 45 sec.

### Bioinformatics

Phosphopeptides that are more abundant in the two B’ζ mutants than in WT were identified using the DEqMS method [19]. Data was normalized applying equal median normalization before applying DEqMS. Furthermore, unmodified peptides and peptides only oxidized were removed from the dataset. In the comparison between WT, *z1* and *z2*, peptides were considered differentially phosphorylated if *p*_adj_ < 0.05 and absolute log_2_FC > 1. Proteins involved in energy metabolism were subjected to gene expression analysis by Genevestigator [19]. Mass spectrometry-verified phosphorylation sites were searched for in candidate PP2A-B’ζ interactors using the PhosPhAt4 database (https://phosphat.uni-hohenheim.de/) [20].

Subcellular localization predictions were done with the tool TargetP 2.0 (https://services.healthtech.dtu.dk/service.php?TargetP-2.0) [21]. Enriched Gene Ontology (GO) terms [22,23] were identified in the combined phosphoproteome datasets from WT and *z1*. The GO analysis was performed using topGO v. 2.50.0 [24] with the adjusted p-values from the DEqMS test of the WT vs *z1* data. Parameters for the GO analysis test were the default weight01 algorithm and Kolmogorov-Smirnov statistics. Annotations used were from the org.At.tair.db R package [25].

## Results and Discussion

Our previous work showed that B’ζ mutant seedlings displayed a growth retardation phenotype on sucrose-free medium [9], indicating a defect in energy metabolism. In addition, B’ζ was reported to be dually targeted to the cytosol and mitochondrion [7]. We hypothesise that the PP2A-B’ζ holoenzyme is involved in regulation of the mitochondrial succinate/fumarate translocator or affects enzymes involved in the Krebs cycle. To test this, we set out to investigate the potential substrates of PP2A-B’ζ by BiFC and phosphoproteomics analysis.

### Sugar dependence assay points to a role in energy metabolism

To further investigate the function of PP2A-B’ζ, we employed two homozygotic knockout lines (SALK_150586C and SALK_107944C), hereafter referred to as *z1* and *z2*. The knockout genotypes of both lines were confirmed by PCR. Line *z2* was previously observed to have impaired growth on sucrose-free medium [9], whereas line *z1* was added to the present study to strengthen our phenotypic and functional observations. Both mutants displayed impaired hypocotyl elongation on sucrose-free medium, especially when grown under continuous darkness (Fig. 1), clearly indicating a role for the B’ζ subunit in energy metabolism. No phenotype was apparent in *z1* or *z2* when grown on sucrose-containing MS or in soil, which is aligned with observations from Matre et al. [7].

**Figure 1.**
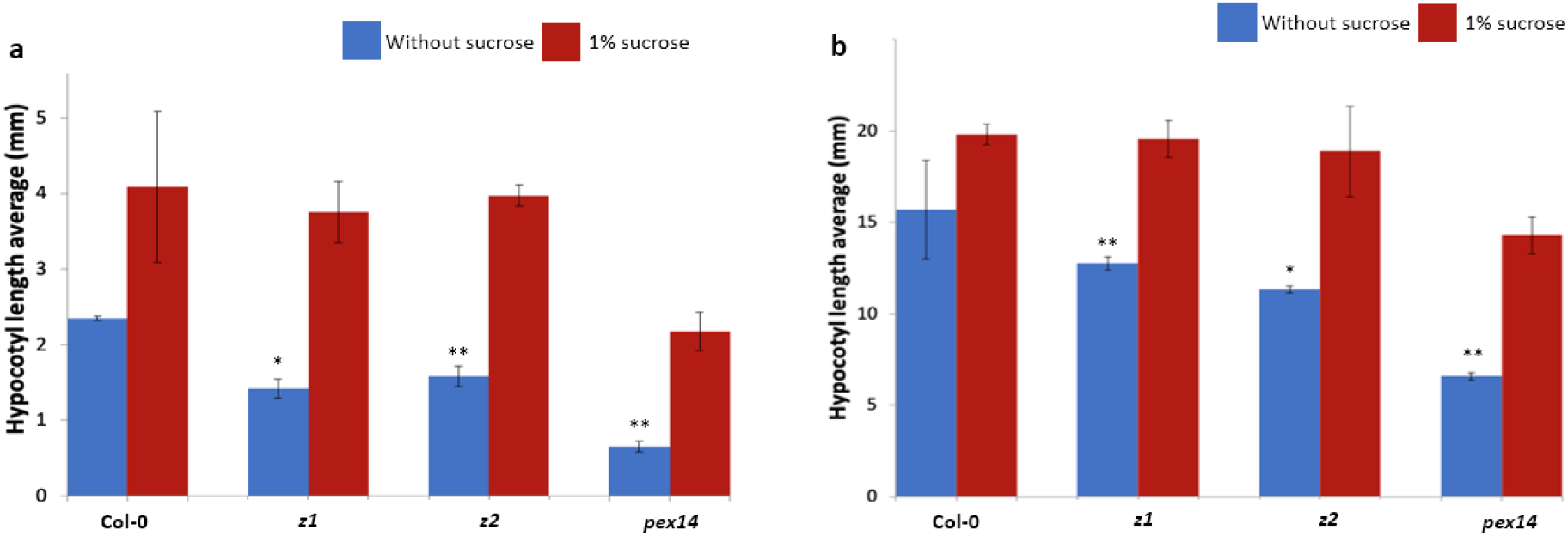
The *pp2a-b’ζ* knockout mutants show reduced growth on medium without sucrose. Hypocotyl length was measured on seedlings grown on MS with or without sucrose in **a** 8 h light/16 h darkness or **b** continuous darkness. Ten roots were measured per genotype and the experiment was repeated four times. * *P* < 0.05 and ** *P* < 0.01; two-tailed t-test with equal variance (no sucrose vs 1% sucrose). Error bars indicate standard deviation.

These findings implicate a putative role for PP2A in regulating the energy metabolism through mitochondria. Therefore, we investigated the phosphor-status of the enzymes of the Krebs cycle, with the aim of identifying potential substrates for PP2A-B’ζ. Seven candidates were targeted, based on their documented key roles in the Krebs cycle: Citrate synthase (CSY5), Aconitase (ACO3), Succinate dehydrogenase (SDH2-1), Mitochondrial succinate/fumarate carrier (mSFC1), Succinyl-CoA-ligase (SCS), Fumarase (FUM1) and Mitochondrial malate dehydrogenase (mMDH2). Since all candidates except mSFC are part of multigene families in *Arabidopsis*, we carefully chose for each protein the paralog that was most relevant for our study. For example, ACO3 carries the main aconitase activity out of the three *Arabidopsis* ACOs [26] and FUM1 was preferred over FUM2 since it is mitochondrion-targeted and is essential (i.e., FUM2 cannot compensate for the lack of FUM1) [27].

### Selected B’ζ putative interactors carry Ser and Thr phosphorylation

First, we analysed the gene expression pattern of *B’ζ* and the genes coding for the seven metabolic proteins using Genevestigator [19]. All genes were found to be expressed throughout development, but the *mSFC1, ACO3* and *B’ζ* genes found to be upregulated under leaf senescence (Supplementary Fig. S2). Next, we looked for evidence for Ser and Thr phosphorylation sites in the candidate substrates using the PhosPhAt4 database [20]. Experimentally verified phospho-Ser and phospho-Thr residues were identified in ACO3, FUM1 and SCS, making them plausible targets for dephosphorylation by PP2A-B’ζ (Supplementary Table S2). On the other hand, mSFC1, SDH2-1, mMDH2 and CSY5 each contain several predicted phosphorylation hotspots (Supplementary Table S2).

### Verification of B’ζ putative interactors involved in energy flow to mitochondria

The genes encoding each enzyme and the carrier protein were cloned and the proteins were tested *in vivo* for their interaction with B’ζ by BiFC, a method that allows detection of protein-protein interactions that under natural conditions are transient and thus hard to observe. Two heterologous systems for plant transient expression were used: tobacco leaves (agroinfiltration) and onion epidermal cells (biolistic bombardment). Since mitochondrial localization of B’ζ depends on a free N-terminus [7], we tagged B**’**ζ with part of the fluorescent protein Venus in its C terminus. The other part of the fluorescent protein was linked to the C terminus of each candidate interactor (Supplementary Fig. S1).

Through transient expression in tobacco leaves, the B’ζ fusion protein was observed to interact with mMDH2 in organelle-like structures, which appears to be the mitochondria (Fig. 2a). This association of B’ζ with a key enzyme of the Krebs cycle provides a first link between PP2A and the mitochondrial energy flow. mMDH2 is one out of two mitochondrial-localized MDH enzymes in *Arabidopsis* and catalyzes the conversion of malate into oxaloacetate in the Krebs cycle. The mMDH enzymes play key roles in the early life stages in *Arabidopsis*, as evidenced from the severe phenotype of the *mmdh1mmdh2* double knockout [28]. We also found a positive interaction of B’ζ with mSFC1 (Fig. 2b). This association takes place in tobacco mesophyll cells, aligning with the reported localization of YFP-tagged B’ζ in *Arabidopsis* [7]. Additionally, fluorescent signal was observed upon the investigation of the interaction between B’ζ and ACO3 in onion epidermal cells (Fig. 3a). In addition, physical interaction between B’ζ and mSFC1 was confirmed (Fig. 3c).

**Figure 2.**
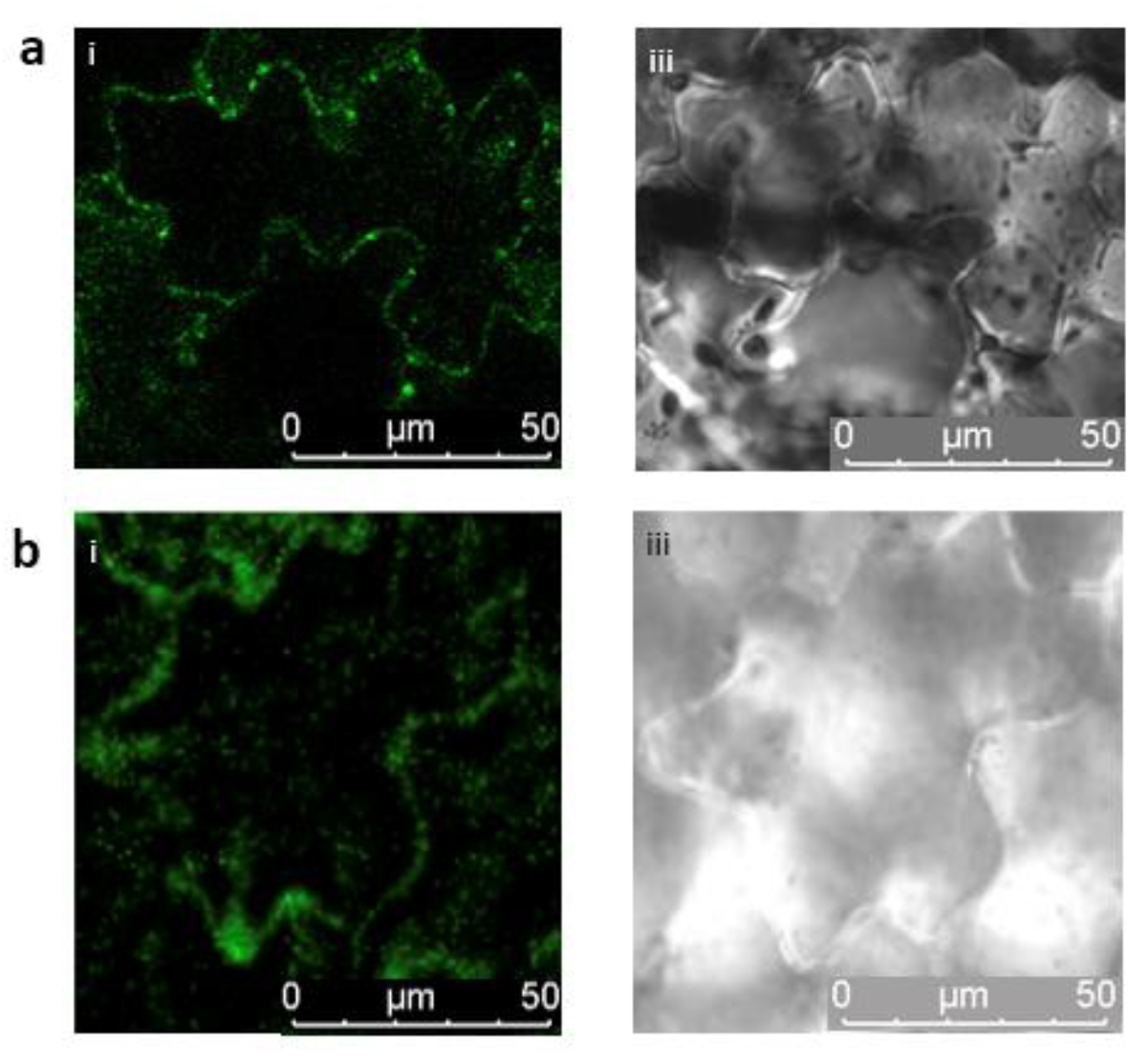
Interaction of PP2A-B’ζ with mMDH2 and mSFC1. BiFC constructs were transiently expressed in tobacco leaf epidermal cells. The C-terminal part of Venus was fused to B’ζ and the N-terminal Venus fragment was fused to **a** mMDH2 and **b** mSFC1. **i** Venus; **ii** brightfield. Both interactions are suggested to be mitochondrial.

**Figure 3.**
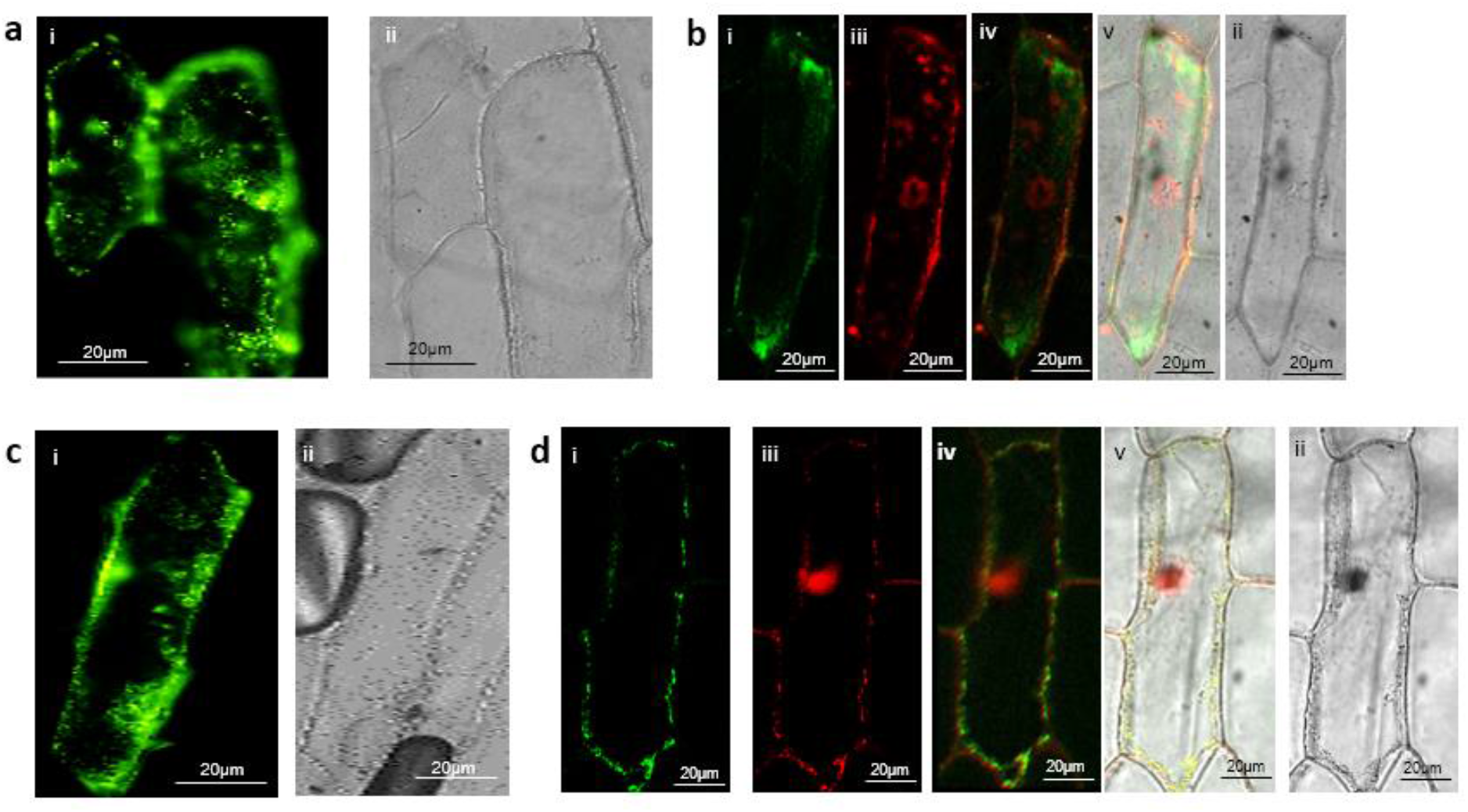
Visualization of PP2A-B’ζ interaction with mSFC1 and ACO3 in onion epidermal cells via BiFC. B’ζ was tagged-terminally with the C-terminal part of Venus while the N-terminal portion of Venus was fused to the C terminus of ACO3 (**a** and **b**) and mSFC1(**c** and **d**). **i** Venus; **ii** brightfield; **iii** RFP (mitochondrial marker pWEN95); **iv** overlay of Venus and RFP; **v** overlay of Venus, RFP and brightfield. In d the signals from Venus and the mitochondrial marker overlap, indicating mitochondrial localization of the mSFC1-B’ζ interaction. The absence of overlap with the mitochondrial marker in b suggests cytoplasmic localisation of the ACO3-B’ζ interaction.

To determine the localization of the positive interactions, the mitochondrial marker pWEN95 was used. The mitochondrial marker harbors the mitochondrial import peptide from the β-subunit of the mitochondrial ATP synthase gene [29], fused with red fluorescent protein (RFP). The import peptide targets RFP to mitochondrial matrix. The interaction complex including the succinate/fumarate carrier, mSFC1, was shown to be colocalized with the mitochondrial marker (Fig. 3d), suggesting that the interaction of PP2A-B’ζ and mSFC1 takes place in mitochondria or on the mitochondrial membrane in onion cells. Due to the mechanism of mitochondrial protein import, which requires protein unfolding for passage through the mitochondrial import system [30], the PP2A holoenzyme would need to disassemble before entry and then reform inside the mitochondrion. A mitochondrial targeting signal in one of the subunits would therefore not be able to direct the PP2A complex into the matrix. More likely, therefore, the interaction between mSFC1 and PP2A takes place on the mitochondrial surface. This interpretation agrees with the previous observation of mSFC1-GFP fusion protein at the mitochondrial membrane in tobacco epidermal cells [31].

The interaction complex including the ACO3, on the other hand, was seen to be cytoplasmic (Fig. 3b). Overall, our BiFC data agree with previously reported localization of B’ζ to both the mitochondrion and the cytoplasm [7]. ACO3 is an iron-sulfur-containing hydratase that is dually targeted to the mitochondrion and the cytoplasm in plants and yeast [26,32,33]. Mitochondrial ACO3 is active in the Krebs cycle, where it catalyzes the stereo-specific isomerization of citrate to isocitrate. The cytosolic form of *Arabidopsis* ACO3 was implicated in the regulation of organellar reactive oxygen species (ROS) homeostasis and was reported to bind to the PP2A-B’γ subunit [34]. Our observation of cytosolic interaction between ACO3 and B’ζ indicates that PP2A-B’ζ could potentially be involved in ROS signalling as well, and that ACO3 could be putatively dephosphorylated by two different PP2A isozymes from the same PP2A subgroup.

Co-localization of B’ζ with mSFC1 and ACO3 is in line with concordant up-regulation of their three respective genes in senescent leaves (Supplementary Fig. S2). No interaction was seen between PP2A-B’ζ and CSY5, SDH2-1, SCS or FUM1 in tobacco and onion cells. Possibly, dephosphorylation of these metabolic enzymes takes place in other developmental stages, in different tissue types or under specific conditions not tested here. In summary, BiFC revealed cytoplasmic interaction of PP2A-B’ζ with the metabolic enzyme ACO3 and the transporter protein mSFC1. The latter most likely takes place at the mitochondrial surface. All observations were strengthened from multiple independent transformation events and in two heterologous plant expression systems.

The mSFC1 carrier protein plays an important role in the exchange of metabolites between organelles [35]. By inference from the homologous protein in yeast, *Arabidopsis* mSFC1 was first believed to import peroxisomal-produced succinate into the mitochondrion in exchange for fumarate, which in turn via multiple downstream reactions would be used in gluconeogenesis or in the glyoxylate cycle [27,35]. However, in the absence of biochemical evidence for succinate/fumarase exchange of mSFC1, its function remained obscure, until more recently, it was revealed that mSFC1 transports citrate, isocitrate and aconitate more efficiently than fumarate and succinate. Thus, the main role of mSFC1 was proposed to be the exchange of cytosolic citrate for mitochondrial isocitrate [31]. The mSFC1 was also found to play an important function in seed germination and seedling development, which is consistent with its expression in cotyledons, hypocotyls and root tips [31,35]. Physical interaction between B’ζ and mSFC1 suggests that a putative dephosphorylation mediated by the PP2A-B’ζ complex could play a role in regulation of metabolite exchange and cellular energy production in *Arabidopsis*.

### Additional metabolic enzymes revealed by phosphoproteomics

To further confirm the protein-protein interaction of PP2A-B’ζ with ACO3 and mSFC1, their encoding cDNAs were cloned into protein expression vectors for investigating the *in vitro* interaction employing microscale thermophoresis (MST). Recombinant proteins were produced in *Escherichia coli* and successfully purified, but the purified protein of PP2A-B’ζ precipitated in all purified trials and hampered the planned protein-protein interaction by MST analysis. We therefore sought to employ comparative quantitative phosphoproteomics as an alternative method for studying B’ζ putative interactors. Quantitative phosphoproteomics is a method that can be used to identify and characterise PTM sites in proteins [36]. When performed on samples collected from suitable mutants, it can reveal novel substrates of kinases and phosphatases. Performing tandem mass tag (TMT) labeling prior to fragmentation increases throughput by enabling multiplex quantification of relative differences between samples. In a typical phosphoproteomics workflow, proteins are first fragmented into peptides by trypsin digestion, then separated by liquid chromatography (LC) and finally quantified by mass spectrometry (MS). Phosphorylated sites are identified using bioinformatic methods applied to the MS data. In order to identify substrates affected by the absence of B’ζ, total proteins were extracted from seedlings of WT and the two knockout mutants, *z1* and *z2*, trypsin digested, and their phosphoproteomes were enriched by TiO2 and detected. The three phosphoproteomes were compared. Although most phosphorylation events are transient, we hoped to identify expected accumulation of phosphorylated ACO3 and mSFC1 and other mitochondrial-related proteins in the knockout mutants of PP2A-B’ζ.

Differential expression analysis (DEqMS) on the phosphoproteomics data was employed. Although, both ACO3 and mSFC1 were not differentiated in the mutants, the phosphorylated peptide form of two mitochondrial proteins involved in plant energy metabolism were found differentially upregulated in one of the knockout mutants (Fig. 4,): Phosphoenolpyruvate carboxylase 3 (PPC3) and Phosphoenolpyruvate carboxykinase 1 (PCK1). These two enzymes, plus a third protein, Pentatricopeptide repeat protein 6 (PPR6), had increased phosphorylation in the *z1* mutant compared to WT (Supplementary Table S3, Supplementary Table S4). All of which implicate a role for PP2A-B’ζ in the mitochondrial metabolism and putatively increase possible substrates of PP2A-B’ζ as expected from abolished PP2A-B’ζ function in the *pp2a-b’ζ* knockout.

**Figure 4.**
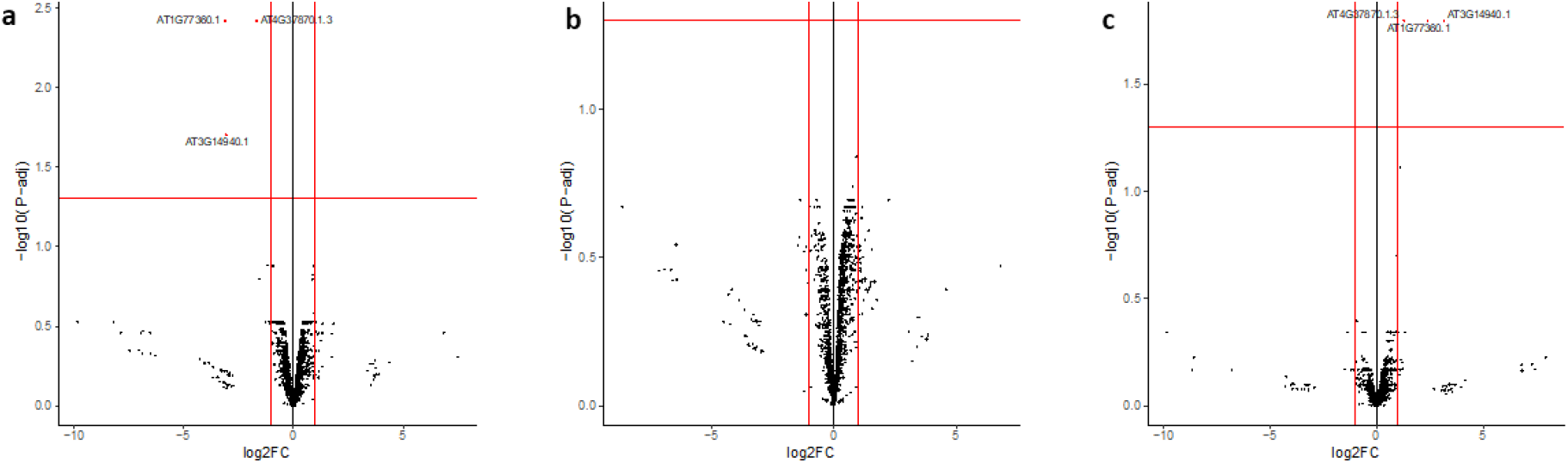
Volcano plots of quantitative phosphoproteomics. Phosphosites with fold-changes >2 and adjusted *p* value <0.05 were regarded as significanlty differentially phosphorylated. The AGI identifiers corresponding to the corresponding three proteins are shown. **a**: WT vs. *z1*. **b**: WT vs. *z2*. **c**: *z1* vs. *z2*.

PPC3 catalyses the carboxylation of phosphoenolpyruvate (PEP) into oxaloacetate (OAA), whereas PCK1 acts in the opposite direction (i.e., forms PEP by OAA decarboxylation). PPC3 and PCK1 thereby play an important role in the regulation of the Krebs cycle. In fact, PCK1 serves as a cataplerotic enzyme, i.e., it removes intermediate metabolites from the Krebs cycle [37]. Interestingly, the two enzymes are known to be regulated by phosphorylation, and PP2A was found to be the phosphatase responsible for dephosphorylating both PPC3 and PCK1 [38,39,40]. The identity of the regulatory subunit was, however, not known in these previous studies. Meanwhile, the Ser and Thr sites that are specifically more abundantly phosphorylated in the *z1* mutant are supported by MS data deposited in the PhosPhAt4 repository (Supplementary Table S2). The detected phosphorylation in PPC3 (Ser 11) is also in agreement with a previous reporting PP2A-medited dephosphorylation at an N-terminal Ser [41,42].

PPR proteins play important roles in post-transcriptional regulation of mitochondrial genes [43,44]. They regulate many different aspects of the RNA life cycle, such as RNA stability, translation, processing, editing and splicing. As such, PPR proteins are important for cellular respiration in plants. PPR6 is predicted to contain a mitochondrial target peptide (TargetP 2.0, likelihood 0.9365), and therefore likely exerts its function inside mitochondria. The maize homolog of PPR6 is responsible for translation initiation and 5’ end maturation of the mitochondrial ribosomal protein S3 (*rps3*) mRNA, with reduced translation and extended *rps3* mRNA 5’ end in the *mppr6* mutant [43]. Since *Arabidopsis PPR6* can complement the maize phenotype, the function of PPR6 seems to be conserved. We therefore hypothesize that PP2A-B’ζ, via dephosphorylation of the PPR6 protein, affects mitochondrial mRNA stability, with consequences for mitochondrial energy metabolism.

Taken together, we supply strong evidence for putative substrates of the PP2A-B’ζ holoenzyme in mitochondria. ACO3 and mSFC1 were not found to be differentially phosphorylated in our comparison between the *z1, z2* and WT phosphoproteomes. Potentially, PP2A-B’ζ shares these two substrates with PP2A holoenzymes containing other regulatory B subunits, which would dephosphorylate ACO3 and mSFC1 in the absence of B’ζ. The difference between our BiFC observations and phosphoproteomics data could also be due to the different tissue types being used – leaves from adult tobacco plants and onion epidermal cells vs. 6-day old whole *Arabidopsis* seedlings.

To get additional insights into which pathways and processes might be regulated by protein phosphorylation-dephosphorylation processes, a GO term enrichment analysis was done. Since the number of differentially phosphorylated proteins in the *z1* mutant was rather low, the combined phosphoproteomic datasets from WT and *z1* were included in the pathway analysis. Notably, the terms phosphoenolpyruvate carboxylase activity (GO:0008964) and phosphoenolpyruvate carboxykinase activity (GO:0004612) were among the most significantly enriched terms in the Molecular Function domain (Supplementary Table S5), supporting the important role of reversible phosphorylation in PEP ← → OAA conversion and the regulation of energy metabolism.

In summary, our phosphoproteomics analysis revealed for the first time that the PP2A-B’ζ subunit could be dephosphorylating the two metabolic enzymes PPC3 and PCK1, in addition to the mitochondrial gene regulator PPR6. All of which could be linked to the role of PP2A-B’ζ holoenzyme in plant innate immunity and energy metabolism that are related to the reported phenotypes in their knockout mutants.

## Conclusion

Knockout mutants of *PP2A-B’ζ* display impaired growth upon sucrose starvation, suggesting a deficiency in energy metabolism when the B’ζ regulatory subunit is removed. In that regard, the TCA cycle enzymes PCK1 and PPC3, as well as the proposed mitochondrial gene expression regulator PPR6 were found differentially phosphorylated in the *pp2a-b’ζ1* mutant. Interaction between PP2A-B’ζ and the metabolic enzymes mMDH2 and ACO3 was confirmed *in vivo*. In addition, the mitochondrial metabolite carrier mSFC1 was shown to interact with B’ζ on the mitochondrial membrane. Taken together, results from this work (Fig. 5) suggest that the PP2A-B’ζ holoenzyme dephosphorylates several proteins involved in mitochondrial energy flow and thus plays an important role in plant metabolism.

**Figure 5.**
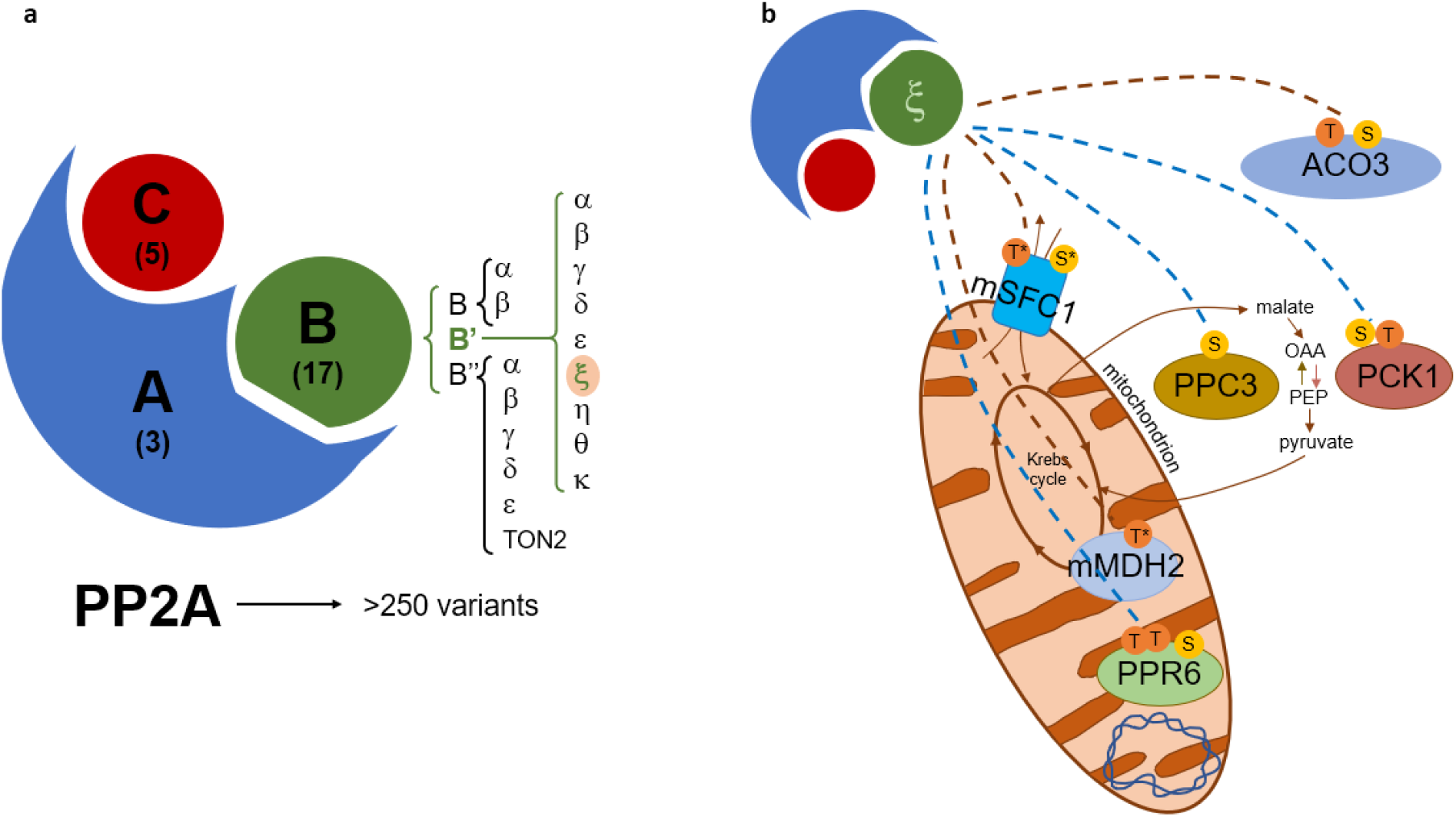
Models of PP2A subunit composition and proposed function of PP2A-B’ζ. **a** The PP2A holoenzyme is composed of a scaffolding subunit (“A”) a regulatory subunit (“B”) and a catalytic subunit (C). The number of genes encoding each subunit in *Arabidopsis* is shown in parenthesis. The focus of this study, ζ, belongs to the B’ protein family. **b** PP2A-B’ζ dephosphorylates multiple metabolic proteins. BiFC-validated substrate interactions of B’ζ are indicated with brown dashed lines and B’ζ -dependent phosphorylation sites revealed by phosophoproteomics are shown with blue dashed lines. Residues known to be phosphorylated are shown in the six substrates. S; serine, T; threonine, *, predicted phosphorylation. PCK1 catalyses the conversion of oxaloacetate (OAA) into phosphoenolpyruvate (PEP), whereas PPC3 catalyses the reverse reaction. Dephosphorylation of the mitochondrion-localised substrates could take place before entry into the mitochondrion or at the mitochondrial membrane (as is most likely for the carrier protein mSFC1).

## Author Contributions

AE and ARAK designed the research; AE and ARAK performed the experiments; AE and KH analyzed the data; AE, ARAK, CL and KH wrote the manuscript. All authors have seen and approved the manuscript and its contents and are aware of the responsibilities connected to authorship.

## Data availability

Unprocessed LC-MS/MS phosphosite data generated in this study is attached (supplementary dataset.xlsb). Corresponding raw files are available upon request.

