## Supplementary material for "Protein-Protein interaction and quantitative phosphoproteomic studies reveal potential mitochondrial substrates of protein phosphatase 2A-B’ζ holoenzyme": Suppl_v3 .pdf

### Identification of novel interactors of PP2A-B'ζ

**Fig. S1** Schematic representation of BiFC constructs.
**Fig. S2** GeneVestigator expression study of genes coding for proteins involved in energy metabolism.
**Table S1** Oligonucleotide sequences for genotyping and cloning.
**Table S2** Phosphopeptides retrieved from PhosPhAt4 and the literature.
**Table S3** PP2A-B'Z interactors identified by phosphoproteomics.
**Table S4** Output from phosphoproteomics DEqMS analysis.
**Table S5** Output from Gene Ontology analysis.

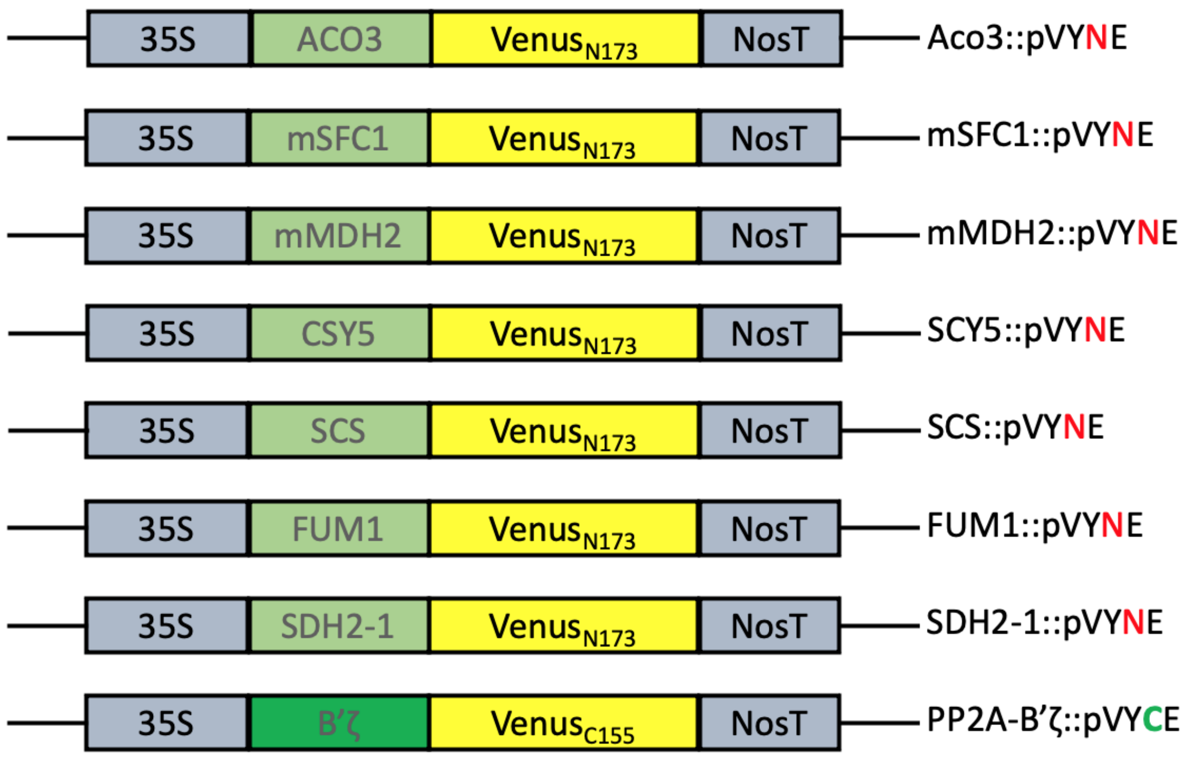


**Supplementary Fig. S1** Schematic representation of BiFC constructs. The seven interactor candidates were fused at their respective C-terminus to the N-terminal part of Venus (pVYNE vector), while the regulatory subunit PP2A-B'ζ was fused at its C-terminus to the C-terminal part of Venus (pVYCE vector). All fusion proteins were expressed from the CaMV 35S promoter. N173: amino acids 1-173 of Venus. C155: Venus amino acids 155-238 (Shyu et al., 2006).

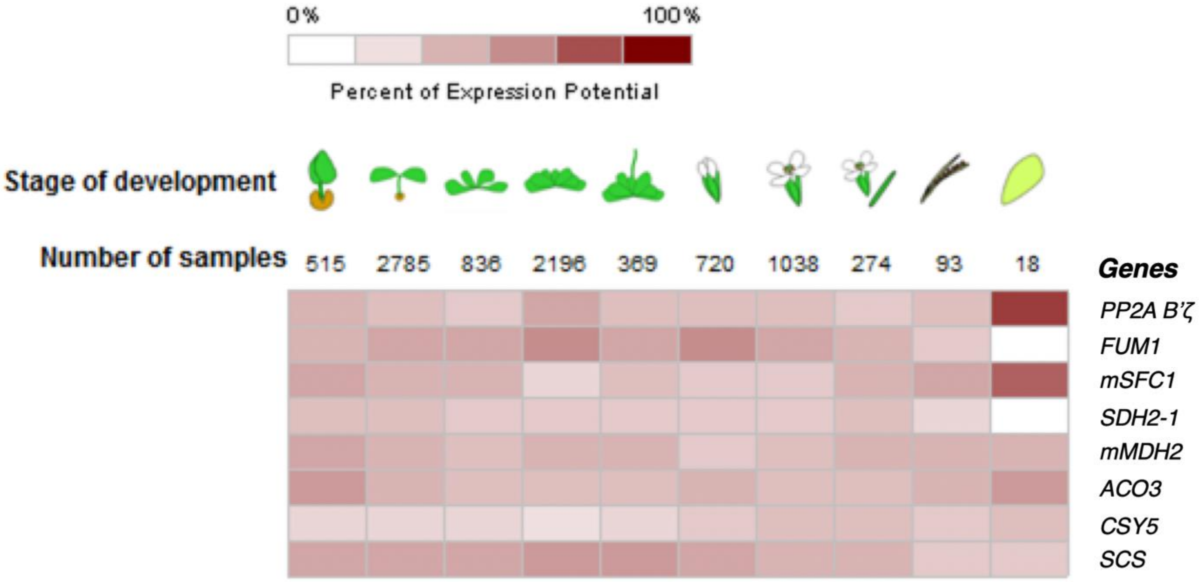


**Supplementary Fig. S2** GeneVestigator expression study of genes coding for proteins involved in energy metabolism. All genes are expressed throughout development, and *PP2A-B'ζ*, *mSFC1* and *ACO3* are upregulated in senescent leaves.

**Table S1. Oligonucleotide sequences for genotyping and cloning.**

| Name | Template | Sequence | Usage |
|---|---|---|---|
| Z1_150586_LP | AT3G21650 | TTTTCACTTCAGAGTCAGCCG | Genotyping |
| Z1_150586_RP | AT3G21650 | ATGGTGCATCGACCTTACATC | |
| Z2_107944_LP | AT3G21650 | CACTCGTCGAAAAGAACTTGG | |
| Z2_107944_RP | AT3G21650 | CCGAATCTCTTTATCGGGAAG | |
| Z_F | AT3G21650 | ATAGTCGACATGATCAAACAGATATTTGG | Cloning (BiFC Venus) |
| Z_R | AT3G21650 | ATGGTACCCGACCCTGTGGACTCAGA | |
| FUM1_F | AT2G47510 | ATACTAGTATGTCGATTTACGTCGCGTCG | |
| FUM1_R | AT2G47510 | ATATCTCGAGATCGGAGGGACCAATCAT | |
| mSFC1_F | AT5G01340 | ATACTAGTATGGCGACGAGAACGGAA | |
| mSFC1_R | AT5G01340 | ATATCTCGAGTAAAGGAGCATTCCGAAG | |
| SDH2-1_F | AT3G27380 | ATACTAGTATGGCGTCTGGTTTGATCG | |
| SDH2-1_R | AT3G27380 | ATATCTCGAGACGCTGAAGTTGCTTGAT | |
| mMDH2_F | AT3G15020 | ATACTAGTATGTTCCGATCAATGATTGTTC | |
| mMDH2_R | AT3G15020 | ATATCTCGAGTTGGTTGGCAAATTTGAT | |
| ACO3_F | AT2G05710 | ATACTAGTATGTATTTAACCGCTTCATCTTCC | |
| ACO3_R | AT2G05710 | ATATCTCGAGTTGCTTGCTCAAGTTTCT | |
| CSY5_F | AT3G60100 | ATACTAGTATGGTGTTTTTTCGCAGCGTAT | |
| CSY5_R | AT3G60100 | ATATCTCGAGGCGGTTCAAGCGCGTGAAGTT | |
| SCS_F | AT5G08300 | ATACTAGTATGTCTAGACAAGTGGCAAGGC | |
| SCS_R | AT5G08300 | ATATCTCGAGCTGCTTCAAAAGACCTCT | |

**Table S2. Phosphopeptides retrieved from PhosPhAt4 and the literature.** Sites are experimentally verified, except if denoted as 'Predicted' on column 5.

| AGI | Annotation | Phosphopeptide | Hits | Reference |
|---|---|---|---|---|
| AT3G14940 | PPC3 | $^{9}$MA**S**IDAQLR$^{17}$ (*) | 34 | Wu et al., 2014; Wu et al., 2013; Menz et al., 2016 |
| AT4G37870 | PCK1 | $^{62}$**S**AP**T**TPINQNAAAAFAAVSEEER$^{84}$ (*)<br>$^{62}$**S**APTTPINQNAAAAFAAVSEEER$^{84}$<br>$^{62}$SAP**T**TPINQNAAAAFAAVSEEER$^{84}$<br>$^{60}$KR**S**AP**T**TPINQNAAAAFAAVSEEER$^{84}$<br>$^{60}$KR**S**APT**T**PINQNAAAAFAAVSEEER$^{84}$<br><br>$^{60}$KR**S**APTTPINQNAAAAFAAVSEEER$^{84}$<br>$^{60}$KRSAP**TT**PINQNAAAAFAAVSEEER$^{84}$ | 1<br>5<br>1<br>1<br>2<br><br>7<br>1 | Zhang et al., 2013<br>Wang et al., 2013<br>Rayapuram et al., 2018<br>Rayapuram et al., 2018<br>Nakagami et al., 2010; Rayapuram et al., 2018<br>Rayapuram et al., 2018<br>Wang et al., 2013 |
| AT1G77360 | PPR6 | MKNV**Y**RVLK | 2 | |
| AT2G05710 | ACO3 | $^{801}$DFN**S**YGSR$^{808}$<br>$^{801}$DFNSYG**S**RR$^{809}$<br>$^{89}$**T**F**S**SMASEHPFK$^{100}$<br>$^{89}$TF**S**SMASEHPFK$^{100}$ | 1<br>1<br>2<br>46 | Roitinger et al., 2015.<br>Roitinger et al., 2015.<br>Rayapuram et al., 2018<br>Rayapuram et al., 2018 |
| AT5G01340 | mSFC1 | $^{94}$QTAFKDSE**T**GUVSNRGRFLSGFG$^{116}$<br>$^{235}$KTRLMAQSRD**S**EGGIRY$^{251}$ | | Predicted (§)<br>Predicted (§) |
| AT2G47510 | FUM1 | $^{194}$TLHSTLESK**S**FEFK$^{207}$ | 1 | |
| AT3G27380 | SDH2-1 | $^{40}$**SS**GGGRGSNLKTFQIYR$^{56}$<br>$^{156}$NQYKSIEPWLKRK**T**PASVPA$^{175}$<br>$^{220}$LLHANRWISD**S**RDE**YT**KERLE$^{240}$ | | Predicted (§)<br>Predicted (§)<br>Predicted (§) |
| AT3G15020 | mMDH2 | $^{174}$VTTLDVVRAR**T**FYAGKSD$^{191}$ | | Predicted (§) |
| AT3G60100 | CSY5 | $^{1}$MVFFRSVSAISRLR**S**RAVQQSSLSNSVRWLHSSE$^{34}$ | | Predicted (§) |
| AT5G08300 | SCS | $^{43}$A**S**DPHPPAAVFVDK$^{56}$<br>$^{185}$IGIMPG**Y**IHKPGK$^{197}$<br>$^{296}$MGHAGAIV**S**GGK$^{407}$<br><br><br>$^{308}$G**T**AQDKIK$^{315}$<br>$^{331}$IGSAM**Y**ELFQER$^{342}$ | 1<br>1<br>4<br><br><br>1<br>5 | Al-Momani et al., 2018<br><br>Nakagami et al., 2010; Rayapuram et al., 2018; Sugiyama et al., 2008<br>Roitinger et al., 2015<br>Wu et al., 2013 |

(*) Identical to phosphopeptide detected in our phosphoproteomics analysis (Table S3).
(§) Phosphorylation site hotspot predicted by PhosPhAt4 (https://phosphat.uni-hohenheim.de/).

**Table S3. PP2A-B'Z interactors identified by phosphoproteomics.**

| AGI | Annotation | Detected phosphopeptide | Probability |
|---|---|---|---|
| AT3G14940 | PPC3 | $^{9}$MA**S**IDAQLR$^{17}$ | 1 |
| AT4G37870 | PCK1 | $^{62}$**S**AP**T**TPINQNAAAAFAAVSEEER$^{84}$ | 0.967, 0.776 |
| AT1G77360 | PPR6 | $^{461}$**TT**QKACVLLEEMIEMGIRP**S**GVTFGR$^{486}$ | 1, 1, 0.786 |
